# Improving recovery of member genomes from enrichment reactor microbial communities using MinION–based long read metagenomics

**DOI:** 10.1101/465328

**Authors:** Krithika Arumugam, Irina Bessarab, Xianghui Liu, Gayathri Natarajan, Daniela I. Drautz–Moses, Stefan Wuertz, Federico M. Lauro, Ying Yu Law, Daniel H. Huson, Rohan B. H. Williams

## Abstract

New long read sequencing technologies offer huge potential for effective recovery of complete, closed genomes. While much progress has been made on cultured isolates, the ability of these methods to recover genomes of member taxa in complex microbial communities is less clear. Here we examine the ability of long read data to recover genomes from enrichment reactor metagenomes. Such modified communities offer a moderate level of complexity compared to the source communities and so are realistic, yet tractable, systems to use for this problem. We sampled an enrichment bioreactor designed to target anaerobic ammonium-oxidising bacteria (AnAOB) and sequenced genomic DNA using both short read (Illumina 301bp PE) and long read data (MinION Mk1B) from the same extraction aliquot. The community contained 23 members, of which 16 had genome bins defined from an assembly of the short read data. Two distinct AnAOB species from genus *Candidatus* Brocadia were present and had complete genomes, of which one was the most abundant member species in the community. We can recover a 4Mb genome, in 2 contigs, of long read assembled sequence that is unambiguously associated with the most abundant AnAOB member genome. We conclude that obtaining near closed, complete genomes of members of low-medium microbial communities using MinION long read sequence is feasible.

---

The development of long read sequencing technologies, such as the Oxford Nanopore Technology MinION and Pacific Biosciences SMRT offer huge potential for effective recovery of complete, closed genomes [1, 2]. While much progress has been made towards this objective for single species isolates [3, 4], the ability of these methods to recover genomes of member taxa in complex microbial communities (microbiomes) is less clear.

To date there is limited long read data available in the metagenome context. Several recent studies have used MinION or PacBio sequencing on highly enriched reactor communities, for example a recent study using PacBio long read sequencing to recover a complete, closed genome of *Kuenenia stuttgartiensis* which was present at around 95% relative abundance [5]. Other studies have obtained long read data or combined long and short read data on more complex communities, for example from moderately enriched bioreactor communities [6], co-culture enrichments [7], marine holobionts [8] or from full scale anaerobic digester communities [9]. There are also several datasets which provide long and short read data from mock communities [10, 11, 12], and which constitute important new bench-marking data. New long read analysis methods [13] and binning algorithms designed for long read metagenome data [14] are becoming available, anticipating the future expansion of metagenome data from these new instruments.

Here we examine the ability of long read data to recover genomes from enrichment bioreactor metagenomes. Such modified communities offer a moderate level of complexity compared to the source communities and so are realistic, yet tractable, systems to use for this problem. Specifically, we sampled an enrichment bioreactor targeting anaerobic ammonium-oxidising bacteria (AnAOB), extracting genomic DNA and obtaining both short read (Illumina MiSeq 301bp PE) and long read data (MinION Mk1B) from the same aliquot. We perform a comparative analysis of different long read assemblers and assembly workflows, using the draft genomes extracted from the short read assembly as an independent references to assess the ability of long read assemblies to capture genome of member species of the community. Our findings show that the long read metagenomics can recover a complete, near–closed genome of the most abundant community member, even in the presence of multiple species of the same genus.

## Methods

### Overall approach

We studied a bioreactor community established to enrich for anaerobic ammonia-oxidizing bacteria (AnAOB), all known species of which belong to the phylum *Planctomycetes.* Previously we used this enrichment reactor community to recover a draft genome of a member of the genus *Candidatus* Brocadia [15]. We sampled the suspended biomass of the reactor, extracted DNA and performed both Illumina and Nanopore DNA sequencing from the same extracted aliquot. Such enrichment reactor communities have the advantage of being only moderately complex relative to their source communities and therefore provide an ideal tradeoff between realistic complexity and tractability in relation to member genome recovery.

We constructed long read assemblies and then used draft genomes obtained from short read assemblies for the purpose of assessment. This approach takes advantage of the fact of our having obtained data from both sequencing modalities using the same DNA aliquot, and leverages current understanding of short read metagenome assembly binning and quality assessment procedures [16, 17]. We tested five different long read assemblers or assembly workflows, namely `Canu` [18], `Miniasm` [19], `SMARTDenovo` [20], `Spectrassembler` [21] and `Unicycler` [22]. We note that while these softwares are not designed or endorsed for metagenome analysis, our experimental design explicitly tested their ability to be used in this setting.

### Enrichment bioreactor and sampling

As previously described [15], we (Y.Y.L, G.N, S.W) developed a protocol to enrich for AnAOB organisms using a sequencing batch reactor seeded with return activated sludge from a water reclamation plant (Public Utilities Board, Singapore). The reactor was fed synthetic wastewater containing ammonium and nitrite and operated at 35 °C. The reactor showed a characteristic reddish–brown biofilm on the walls of the reactor (characteristic of AnAOB bacteria), and we observed simultaneous ammonium and nitrite depletion consistent with the occurrence of Anammox. We confirmed the presence of AnAOB species within the community using FISH, 16S amplicon sequencing and shotgun metagenomics [15]. At the time of sampling for the present study (10/2016) the reactor community had been established for approximately 9 months.

### DNA extraction

DNA was extracted from sampled biomass with the FastDNA^TM^SPIN kit (MP Biomedicals) for Soil, using using 4× bead beating with a FastPrep homogeniser (MP Biomedicals) followed by cleanup with Genomic DNA Clean & Concentrator^TM^–10 (Zymo Research).

### Short read sequencing

Genomic DNA Library preparation was performed using a modified version of the Illumina TruSeq DNA Sample Preparation protocol. We then performed a MiSeq sequencing run with a read length of 301 bp (paired-end). The raw short read data is publicly available from the NCBI via BioProject accession `PRJNA471614` or Sequence Read Archive accession `SRR8168439`.

### Long read sequencing

Library preparation was constructed from 1.7*μ*g of genomic DNA using the SQK–LSK 108 Ligation Sequencing Kit (Oxford Nanopore Technologies). Sequencing was performed on a MinION Mk1B instrument (Oxford Nanopore Technologies) using a SpotON FLO MIN106 flowcell (FAF05082) and R 9.4 chemistry. Data acquisition was performed using MinKNOW version 1.3.30 running on a HP ProDesk 600G2 computer (64–bit, 16Gb RAM, 2Tb SSD HD) running Windows10. The raw long read data is publicly available from the NCBI via BioProject accession `PRJNA471614` or and Sequence Read Archive accession `SRR7169548`.

### Analysis of short read sequence data

The raw FASTQ files were processed using `cutadapt` (version 1.14) [25] using the following arguments: ‐‐`overlap 10 –m 30 –q 20,20 ‐‐quality-base 33`. We initially characterised the community data using `Ribotagger` [26] selecting the V4 region and annotating ribotags to the SILVA database [27]. We calculated relative abundance of each ribotag by dividing their reads counts by the total read count from all ribotags. Single–sample metagenome assembly of the short read sequence was constructed using `Newbler` v2.9. The contigs generated from short read data are hereafter denoted as *short read assembled contigs* (SRAC). We identified 16S genes within contigs using the ‐‐`search16` module of `USEARCH` (version 10.0.240, 64 bit) [28], and annotated them using the SILVA::SINA webserver (using default parameters) [29]. Genes (open reading frames) were predicted from all contigs using `MetaGeneMark` [30] and annotated to RefSeq NR (January 2016 version) [24] using `DIAMOND` v0.8.22.84 (running default parameters) [31]. We identified putative member genomes using `MetaBAT` [32]. For each identified bin we performed taxonomic level analysis using `CheckM` [33] and by applying the *K*–statistic methods of Williams *et al.* [34].

### Analysis of long read sequence data

Basecalling was performed using `Albacore` version 2.2.4. Adaptor trimming was performed using `Porechop` [35] with default settings. We performed taxonomic analysis of long read data using MEGAN–LR [13]. We assembled long read data using each of the following long read assemblers: `Canu` [18], `Miniasm` [19], `SMARTDenovo` [20], `Spectrassembler` [21] and `Unicycler` [22], using default setting unless otherwise stated. In `Canu` there is a required parameter (`genomeSize`) that sets the expected genome length, and which controls the number of reads used by assembler. Because this estimate is essentially meaningless in the metagenome context, we computed multiple assemblies using the following values of `genomeSize`: 3.4 Mbp, 5 Mbp, 12 Mbp, 25 Mbp, 50 Mbp and 120 Mbp. Contigs generated from long read data are hereafter denoted as *long read assembled contigs* (LRAC). The number of reads used in each assembly was estimated by mapping long read to LRAC sequence with `minimap2` [36] and using `samtools-1.6` to calculate the number of aligned reads [37].

### Comparative analysis of long and short read assemblies

We used `BLASTN` (version 2.4.0+) [23] to examine the degree of sequence alignment between LRAC and SRAC sequences. We treated the LRAC as the subject sequences and the SRAC as the query sequences, using default `BLASTN` parameters and without the results outputted in tabular format. To assess the extent to which SRAC sequence aligned to LRAC sequence, we computed ratio of the alignment length to the length of the query (SRAC) sequence (*al*2*ql*).

## Results and Discussion

### Community profiling and member genome recovery using short read data

Community profiling using RiboTagger [2] detected a total of 23 member taxa (defined by 16S V4 region sequence captured in metagenome reads). Of these 7 taxa showed a relative abundance of at least 5% (Fig. 1A). In order of decreasing relative abundance, those seven taxa are *Candidatus* Brocadia (genus), a member of phylum *Armatimonadetes*, an unknown taxa (no kingdom assigned), a member of genus *Denitratisoma*, a second *Ca.* Brocadia (genus) taxon, a member of family *Anaerolineaceae* and another unknown taxon. The short read data was subsequently assembled (Table S1) and automatically binned (see Methods and Fig. 1B), which resulted in 16 bins being defined, of which 4 demonstrated an estimated completeness of > 95% and contamination < 5% (Table S2). Taxonomic analysis of each bin was based on presence of 16S sequence, marker gene analysis [33] and analysis of ORF-level annotations [34]. The results were qualitatively consistent with the 16S analysis, and two species of *Ca.* Brocadia identified (bin 2 and bin 6, with mean base coverage of 150 and 31, respectively), a member of phylum *Armatimonadetes* (bin 1, mean base coverage 180) and a member of phylum *Chloroflexi* (bin 7, mean base coverage, 150) (Fig. 1B, Table S2–S3). We observed that bin 3, comprised of a set of 26 contigs (approximately 201k bp of sequence, with mean base coverage 142) that are strongly associated with *Ca.* Brocadia (Table S2–S3), overlapped in the coverage–GC plane with bin 2 and we therefore subsequently combined those two bins under the assumption that they were spuriously separated by the binning procedure (referred to hereafter as bin 2).

**Figure 1:**
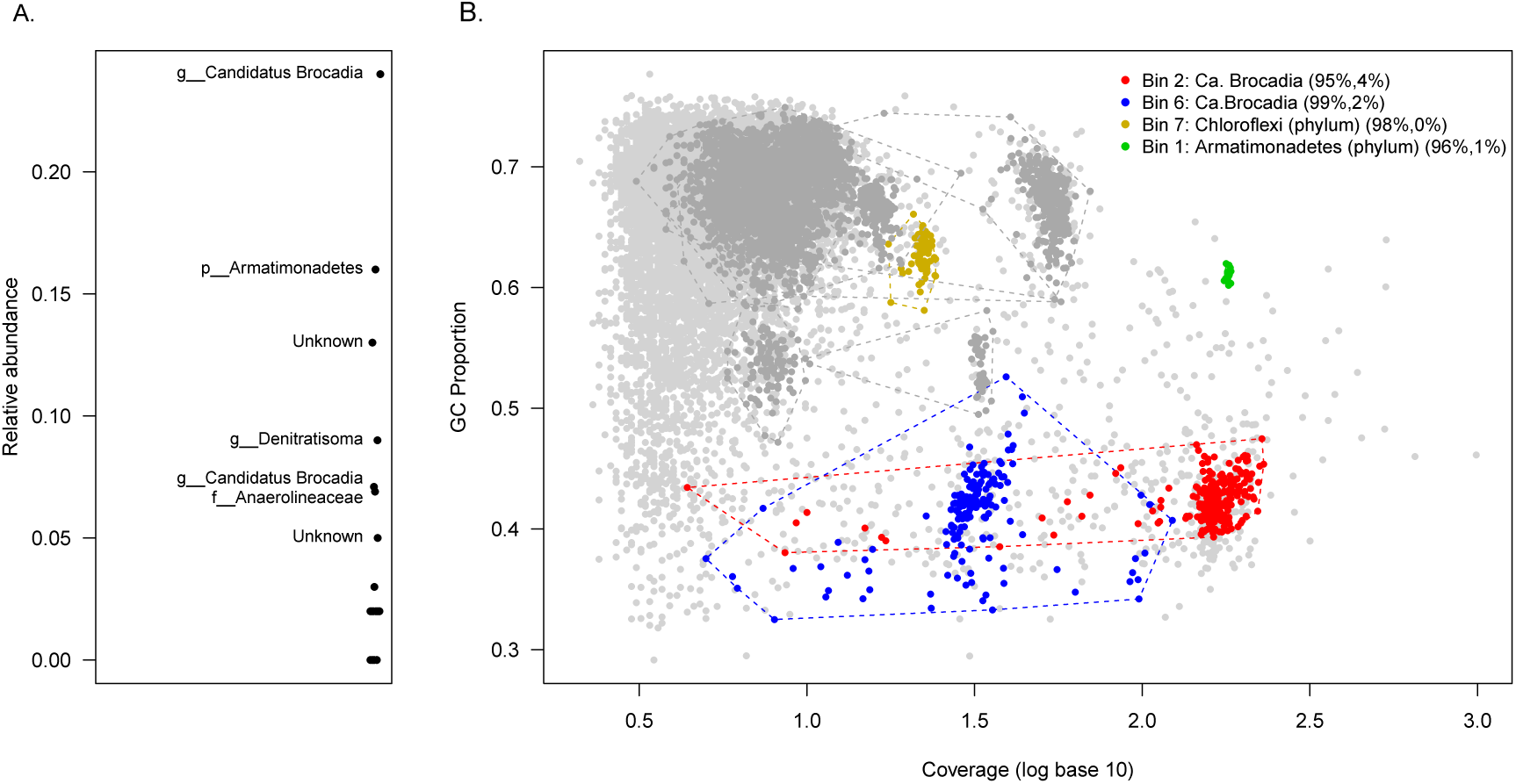
Analysis of short read sequencing data from AnAOB enrichment reactor community. *A*: RiboTagger analysis of short read data, showing taxon ranked by relative abundance. *Unknown* denotes cases in which RiboTagger could not assign any annotation to the corresponding ribotag sequence. A total of 23 taxa were detected, of which 7 had a relative abundance ≥ 0.05 (named). A member of genus *Candidatus* Brocadia, a known AnAOB, was the most abundant community member, with a second member of this genus being present at a lower abundance. *B* : Single–sample metgenome assembly of the short read sequence, constructed using `Newbler` v2.9. GC–proportion and log coverage are used to visualise contigs here. Contigs were binned using `MetaBAT`. A total of 16 bins were detected, of which 4 showed `CheckM` defined completeness ≥ 95% and contamination ≤ 5% (contigs shown in colours and described in legend: numbers in brackets show estimated contamination and completeness). Evidence supporting taxonomic assignments to bin is made on the basis of presence of full length, annotated 16S sequences, marker gene analysis in CheckM and ORF–level annotation statistics (see main text for details).

### Nanopore sequencing and initial taxonomic analysis of long read data

The MinION run generated a total of 115,383 basecalled reads, of which 95,571 held a quality score > 7 (Table 1) and 93.4% were greater than 500 bp in length. The mean read length was 4,677 bp with the longest read being 112,729 bp. There were 6,324 reads with a length greater than 10,000 bp. A total of 539,410,340 bp of sequence was generated from the run.

**Table 1:** Summary statistics for long read data

| Measure |  |
| --- | --- |
| Number of reads | 115,383 |
| Number of reads after adapter trimming | 115,336 |
| Total length of sequenced reads | 539,410,340 |
| Longest read (bp) | 112,729 |
| Shortest read (bp) | 3 |
| Number of reads>500 bp | 107,673 |
| Number of reads>1000 bp | 98,574 |
| Number of reads>10k bp | 6,324 |
| Number of reads>100k bp | 1 |
| Number of reads>1M bp | 0 |
| Mean read length (bp) | 4,677 |
| N50 (bp) | 6,502 |

Taxonomic assignments using MEGAN–LR were obtained for 88,097 reads (76.4% of all reads), representing a total of 326M aligned bases, of which 291Mbp were annotated to bacterial phyla. The most prominent phyla as ranked by aligned base length was Planctomycetes (157Mb or 54% of total sequence aligned to phylum level), *Proteobacteria* (69.7Mb, 24%), *Chloroflexi* (21.0 Mbp) and *Armatimonadetes* (1.79 Mbp, 6.14%) (Table S4). A total of 162 Mbp of alignments were obtained at genus level, with 93.4% of these attributable to *Ca.* Brocadia. Around 2% of aligned bases were attributable to genus *Nitrospira*, with the remaining detected genera each accounting for less than 1% of aligned bases (Table S5).

### Long read assembly

Using the long read data, we constructed assemblies using five assemblers or assembly workflows (Table 2). The total amount of assembled sequence varied between approximately 4.8Mb (`Canu` with `genomeSize` parameter set to 3.4 Mbp) and 11.2 Mbp (`Canu` with `genomeSize` parameter set to 50 Mbp or 120Mbp) (Table 2). Examination of the GC-length data showed a consistent tendency for longest contig to be clustered around a GC of approximately 0.4, consistent with the high abundance of AnAOB (Fig. 2). The distribution of LRAC sequence length varied across assemblers (Fig. 2), but all assemblers returned at least one LRAC sequence greater than 1Mb in length, with `Canu` and `SMARTdenovo` returning 2 sequences longer than 1Mb. Between 33% and 45% of long reads contributed to LRAC sequence, dependent on what assembler or assembly workflow was used (Table 2).

**Table 2:**
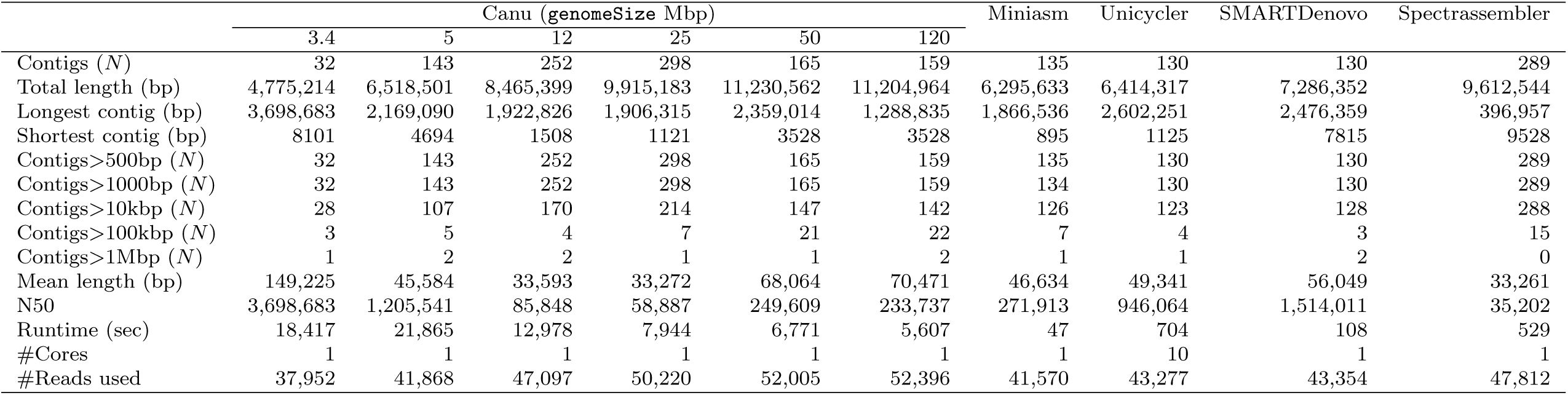
Summary statistics for long read assemblies

**Figure 2:**
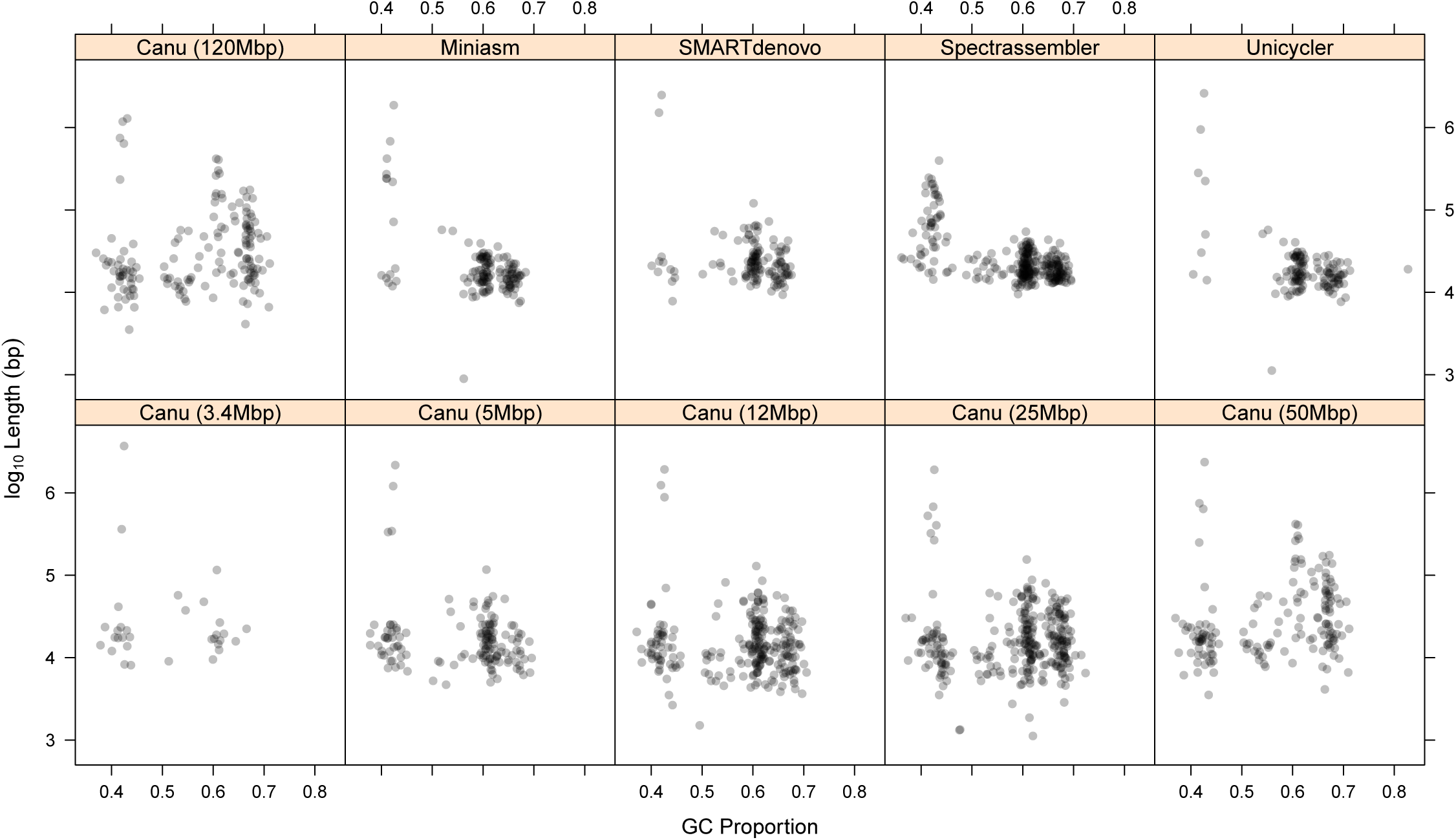
Length distribution of LRAC sequences from each of the five long read assemblers employed in this study

### Taxonomic and genomic analysis of long read assembled sequence

To begin taxomomic analysis of LRAC sequence, we next examined whether 16S gene sequences were present using homology search, and if so, obtained their taxonomic annotations. To do so we aligned all full length 16S genes identified in the short read assembly to all LRAC sequences. In the short read assembly, we identified a full length 16S gene in bin 2 (residing on contig00593) that was aligned to SILVA annotated sequence from *Ca.* Brocadia with 95% identity. That sequence mapped with over 99% identity to LRAC sequences generated by `Canu` and `Unicycler` and close to 99% identity in the case of `Spectrassembler`. In the other two assemblers, the degree of alignment were lower at 88% and 73% for `Miniasm` and `SMARTdenovo`, respectively. A second full length 16S gene annotated to *Ca.* Brocadia in SILVA and recovered from bin 6 was aligned to the same LRAC locations as the bin 2 sequence, but with lower percent identity in all instances (alignment statistics for these two 16S sequences are provided in Supplementary Datafile 1).

We next examined whether these LRAC sequences constituted complete or near–complete genomes of member species. Using BLASTN, we aligned SRAC sequences (treated as queries) to the LRAC sequences (treated as subject) and then categorised the homology search statistics by the bin of origin of the aligned SRAC sequence. We summarised the bulk alignment statistics for each LRAC partitioned by SRAC–bin using the mean of the median of the distribution of the percent identity (*pident*) and the median of the distribution of the alignment length to query (SRAC) length (*al*2*ql*; expressed as a percentage to be on the same scale as *pident*) (Fig. 3). With the exception of `Spectrassembler` the longest LRAC sequences showed a strong degree of alignment with SRAC sequences arising from the highest abundance *Ca.* Brocadia bin (bin 2) and to a lesser degree with SRAC sequences from the lower abundance *Ca.* Brocadia taxon (bin 6). No other LRAC sequence demonstrated a high degree of association with any other genome bin defined in our analysis. We therefore supposed that these LRAC sequences capture substantial sized fragments of the most abundant *Ca.* Brocadia genome present in the community, and then developed further analyses to test this notion. We now focused attention of the `Canu` assembly generated using `genomeSize`=3.4Mb, as this gave the longest single LRAC sequence from any workflow. This LRAC sequence (hereafter referred to as `tig00000001`) contained the full length 16S referred to above. Given this sequence also has a mean GC close to 0.4, we can consider it a candidate for being a large genome fragment of *Ca.* Brocadia present in the reactor community. This contig had a mean long read base coverage of 48 (s.d: 12) (Fig. S1).

**Figure 3:**
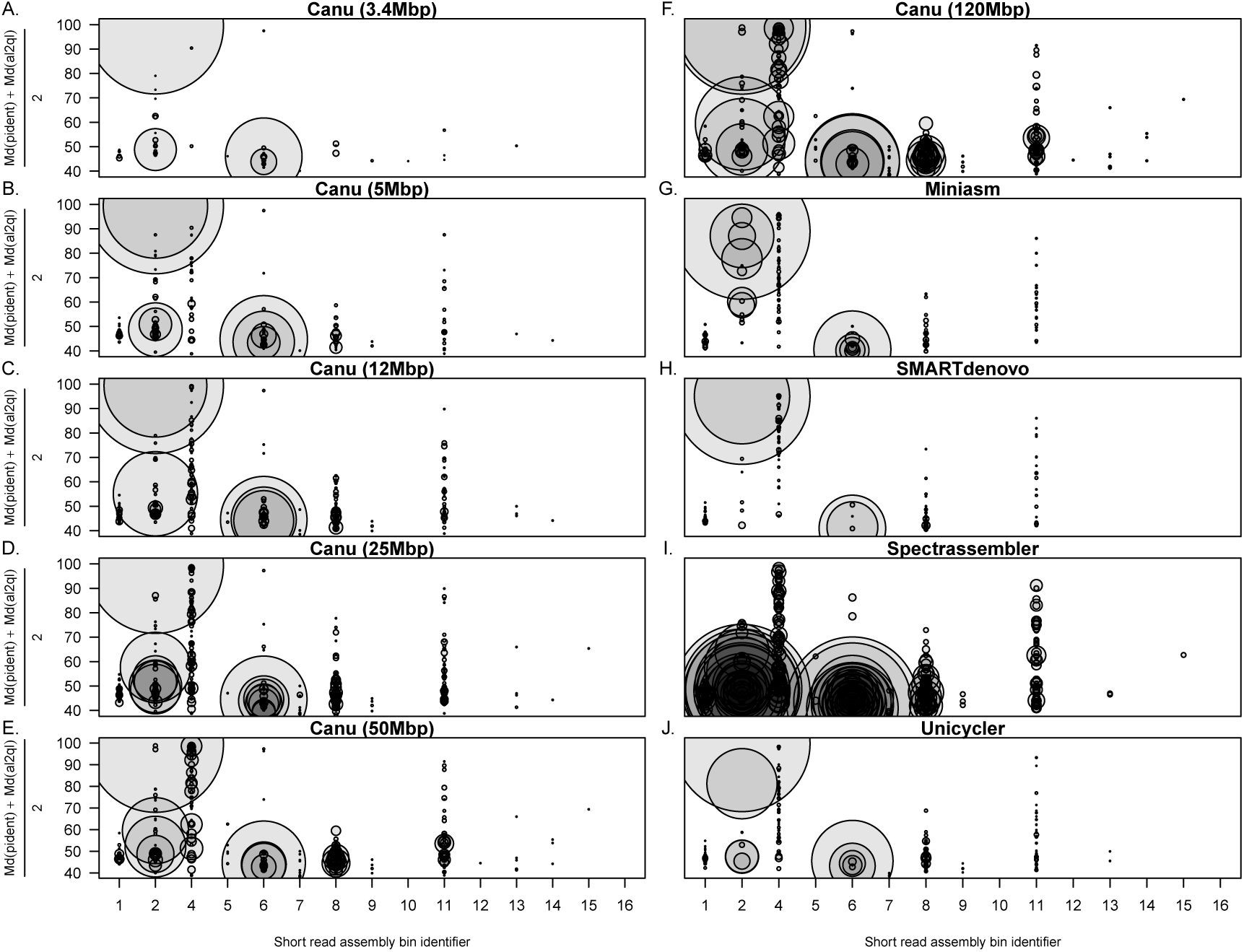
Summary of BLASTN alignment statistics between SRAC sequences and LRAC sequences categorised by bin membership of the SRAC sequences. In each panel, the horizontal axis lists SRAC bin identifiers (see **Figure 1** and **Table S1**). Using BLASTN, SRAC sequences were treated as queries and LRAC sequences as the subject (see **Methods: Comparative analysis of long and short read assemblies**). Each data point corresponds an LRAC sequence with diameter proportional the mean of the median of the distribution of the percent identity (*pident*) and the median of the distribution of the alignment length to query (SRAC) length (*al*2*ql*; expressed as a percentage to be on the same scale as *pident*). Note that the LRAC sequences show a large number of highly aligned SRAC sequences that arising from the *Ca.* Brocadia genome from bin 2, with a lesser degree of alignment evident from the second *Ca.* Brocadia genome captured by SRAC sequence from bin 6.

If this LRAC sequence represents large genomic fragments from a member species of the community, then we would predict the SRAC sequences from the cognate short read assembly bin will align in a spatially uniform fashion across it. We therefore examined the spatial distribution of SRAC alignments across the entire the cohort of LRAC assembly using dot plots, generating these for the following sets of SRAC sequence *1)* all contigs that are members of neither bin 2 nor bin 6; *2)* contigs that are members of bin 6 and *3)* contigs that are members of bin 2. We further subset for those alignments that were full length (defined as alignments exhibiting *al*2*ql* > 95%). The results of each of the corresponding six analyses are shown in Figs 4–9. Each plot shows a set of four panels, the left most of which is a dot plot showing the alignments of the SRAC sequence on the entire LRAC assembly, the next rightmost panels showing the corresponding BLAST statistics for SRAC alignments, namely *al*2*ql* and *pident*, and the rightmost panel shows the length of the aligned SRAC sequences.

**Figure 4:**
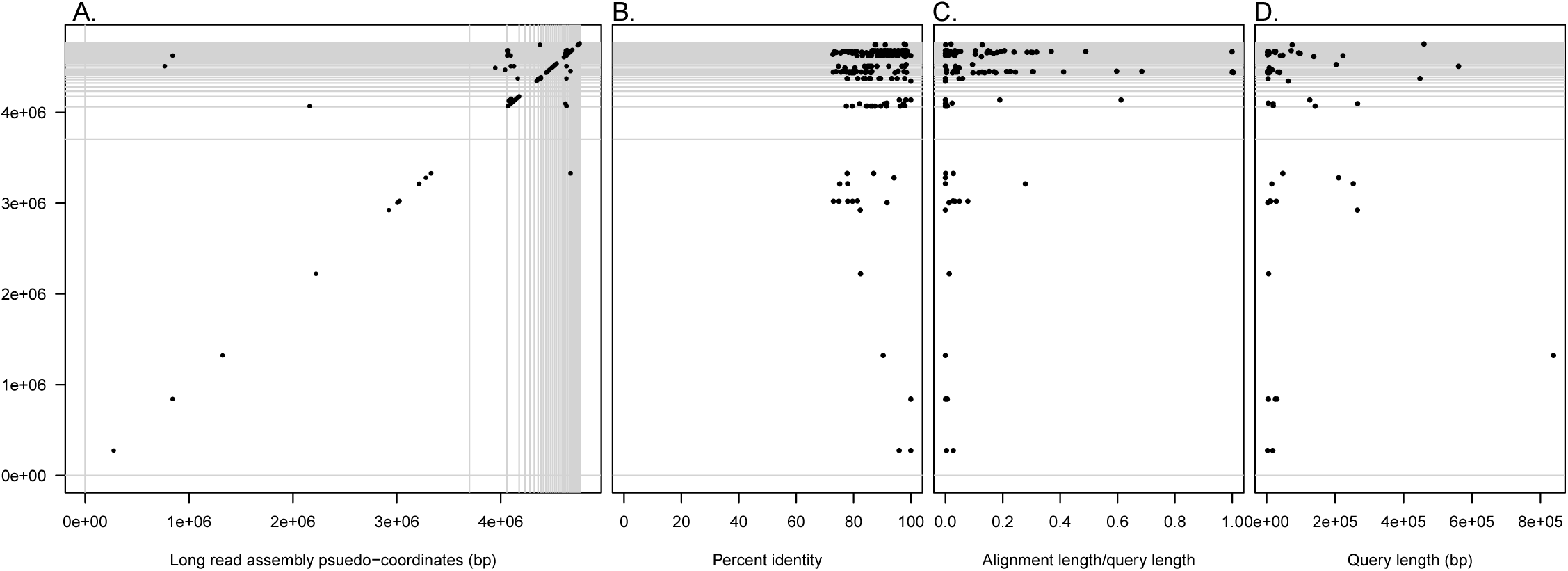
Alignment of SRAC and LRAC sequence for LRAC for SRAC contigs that are **not member of either AnAOB bin**, using **all alignments returned by BLASTN**. *A).* Dot–plot showing LRAC assembly in pseudo–coordinates, ordered by LRAC contig length in increasing order. Vertical and horizontal lines delineate individual contigs. The first, two LRAC contigs are `tig00000001` and `tig00000002`. Black segments denote individual alignments; *B)* percent identity statistics for aligned sequences. Magnitude of percent identity (*pident*) is plotted on *x*–axis with alignment location on the LRAC assembly plotted on the *y*-axis; *C*) query length statistics for aligned sequences. Magnitude of query length is plotted on *x*–axis with alignment location on the LRAC assembly plotted on the *y*-axis.

Starting with the SRAC sequences that are neither members of bin 2 or bin 6, we observe only a small number of alignments to `tig00000001` with low values of *al*2*ql* (Fig. 4), and four alignments of full length, high percent identity to other short LRAC sequences in the assembly (Fig. 5). SRAC sequences from each of the two *Ca.* Brocadia bins showed markedly different patterns of alignment. Only SRAC sequences from bin 2 showed spatially uniform tiling across `tig00000001` (Fig. 8), with a high degree of alignment and percent identity (Fig. 9). SRAC sequences from bin 6 also showing a large number of alignments, but at greatly reduced number when we considered only full length alignments (Fig. 7). Based on this analysis, we can conclude that `tig00000001` appears to derive from genome fragments associated with the *Ca.* Brocadia species captured in bin 2, but not bin 6, in the short read assembly. We note that the reduced degree of percent identity from SRAC sequences from bin 6 is not a consequence low coverage (leading to a systematic reduction in contig length), as a systematic relationship between SRAC length and coverage is only evident at coverage values < 10, not at the coverage ranges observed for either *Ca.* Brocadia bin (Fig. S2A).

**Figure 5:**
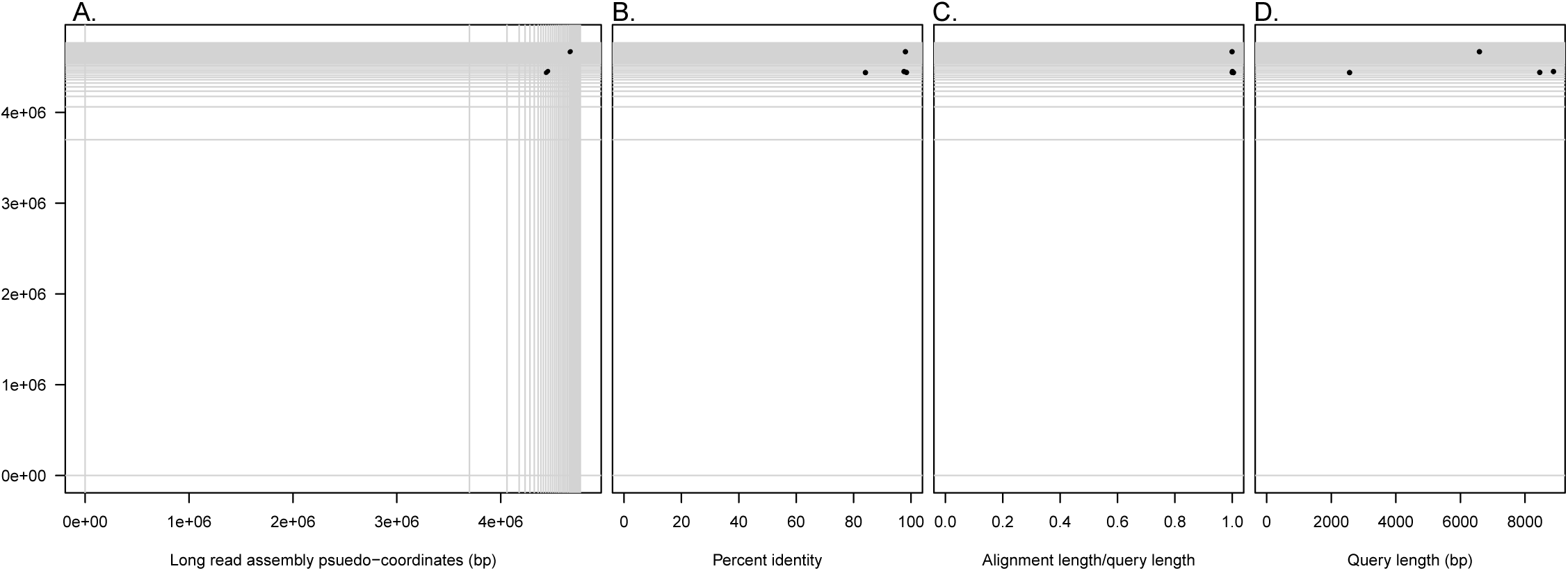
Alignment of SRAC and LRAC sequence for LRAC for SRAC contigs that are **not member of either AnAOB bin**, using **only full length alignments** (*al*2*ql* > 95%). *A).* Dot–plot showing LRAC assembly in pseudo–coordinates, ordered by LRAC contig length in increasing order. Vertical and horizontal lines delineate individual contigs. The first, two LRAC contigs are `tig00000001` and `tig00000002`. Black segments denote individual alignments; *B*) percent identity statistics for aligned sequences. Magnitude of percent identity (*pident*) is plotted on *x*–axis with alignment location on the LRAC assembly plotted on the *y*-axis; *C*) query length statistics for aligned sequences. Magnitude of query length is plotted on *x*–axis with alignment location on the LRAC assembly plotted on the *y*-axis.

**Figure 6:**
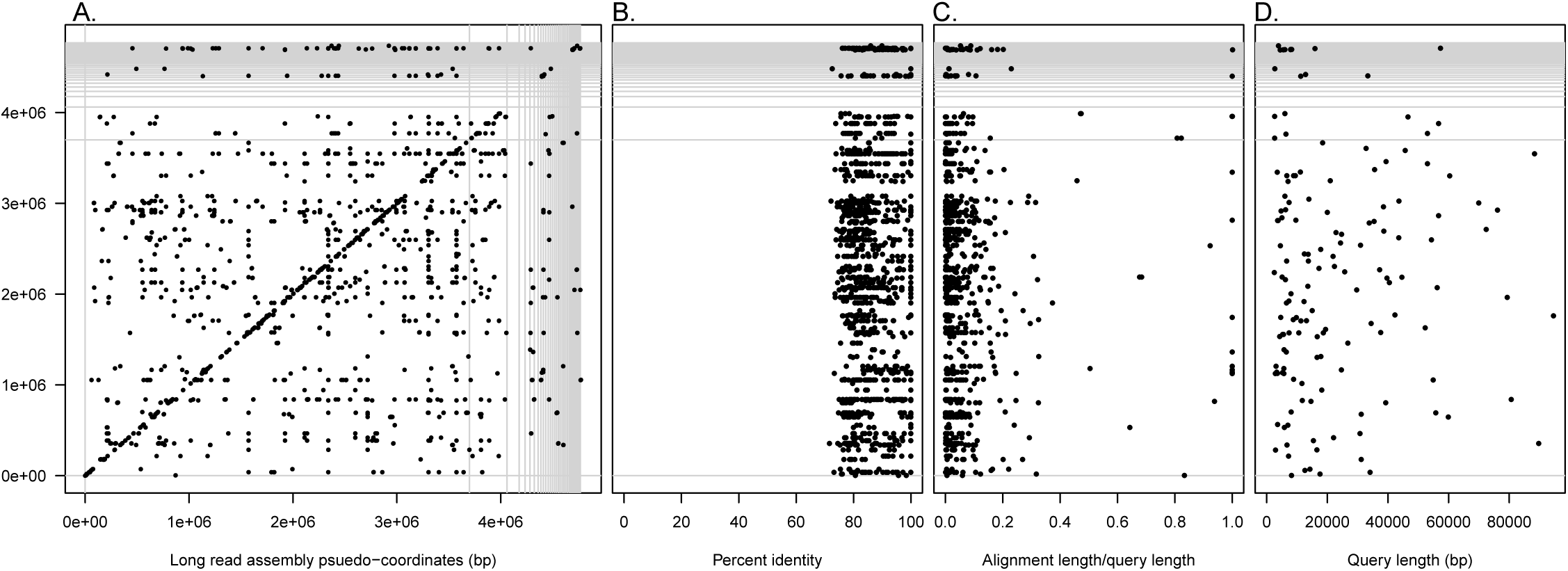
Alignment of SRAC and LRAC sequence for LRAC for SRAC contigs that are **members of AnAOB bin 6**, using **all alignments returned by BLASTN.** *A).* Dot–plot showing LRAC assembly in pseudo–coordinates, ordered by LRAC contig length in increasing order. Vertical and horizontal lines delineate individual contigs. The first, two LRAC contigs are `tig00000001` and `tig00000002`. Black segments denote individual alignments; *B*) percent identity statistics for aligned sequences. Magnitude of percent identity (*pident*) is plotted on *x*–axis with alignment location on the LRAC assembly plotted on the *y*-axis; *C*) query length statistics for aligned sequences. Magnitude of query length is plotted on *x*–axis with alignment location on the LRAC assembly plotted on the *y*-axis.

**Figure 7:**
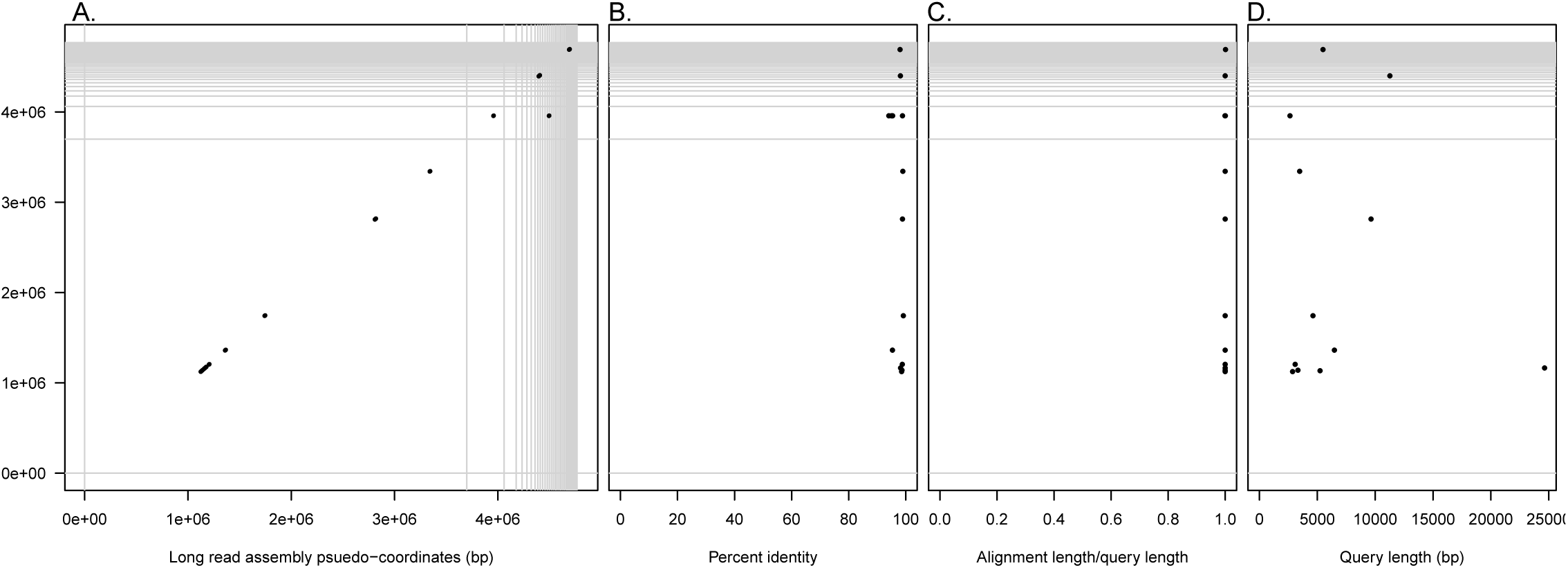
Alignment of SRAC and LRAC sequence for LRAC for SRAC contigs that are **members of AnAOB bin 6**, using **only full length alignments** (*al*2*ql* > 95%). *A).* Dot–plot showing LRAC assembly in pseudo–coordinates, ordered by LRAC contig length in increasing order. Vertical and horizontal lines delineate individual contigs. The first, two LRAC contigs are `tig00000001` and `tig00000002`. Black segments denote individual alignments; *B*) percent identity statistics for aligned sequences. Magnitude of percent identity (*pident*) is plotted on *x*–axis with alignment location on the LRAC assembly plotted on the *y*-axis; *C*) query length statistics for aligned sequences. Magnitude of query length is plotted on *x*–axis with alignment location on the LRAC assembly plotted on the *y*-axis.

**Figure 8:**
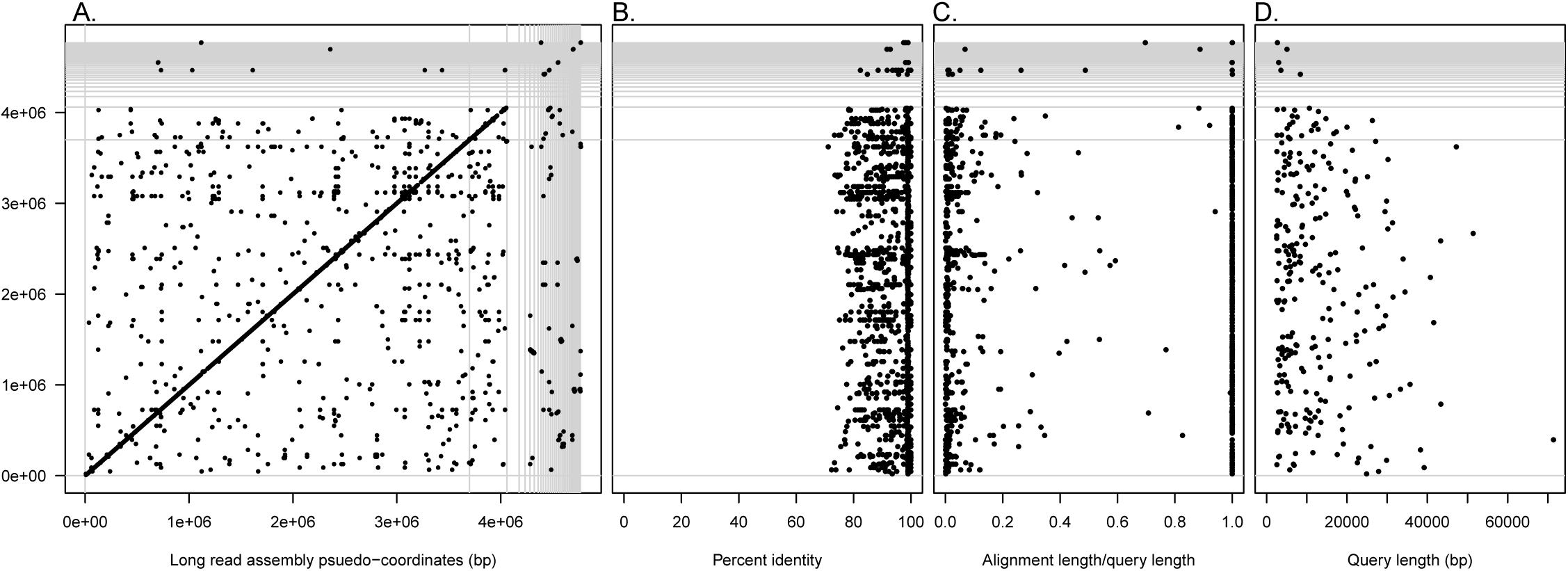
Alignment of SRAC and LRAC sequence for LRAC for SRAC contigs that are **members of AnAOB bin 2**, using **all alignments returned by BLASTN.** *A).* Dot–plot showing LRAC assembly in pseudo–coordinates, ordered by LRAC contig length in increasing order. Vertical and horizontal lines delineate individual contigs. The first, two LRAC contigs are `tig00000001` and `tig00000002`. Black segments denote individual alignments; *B*) percent identity statistics for aligned sequences. Magnitude of percent identity (*pident*) is plotted on *x*–axis with alignment location on the LRAC assembly plotted on the *y*-axis; *C*) query length statistics for aligned sequences. Magnitude of query length is plotted on *x*–axis with alignment location on the LRAC assembly plotted on the *y*-axis.

**Figure 9:**
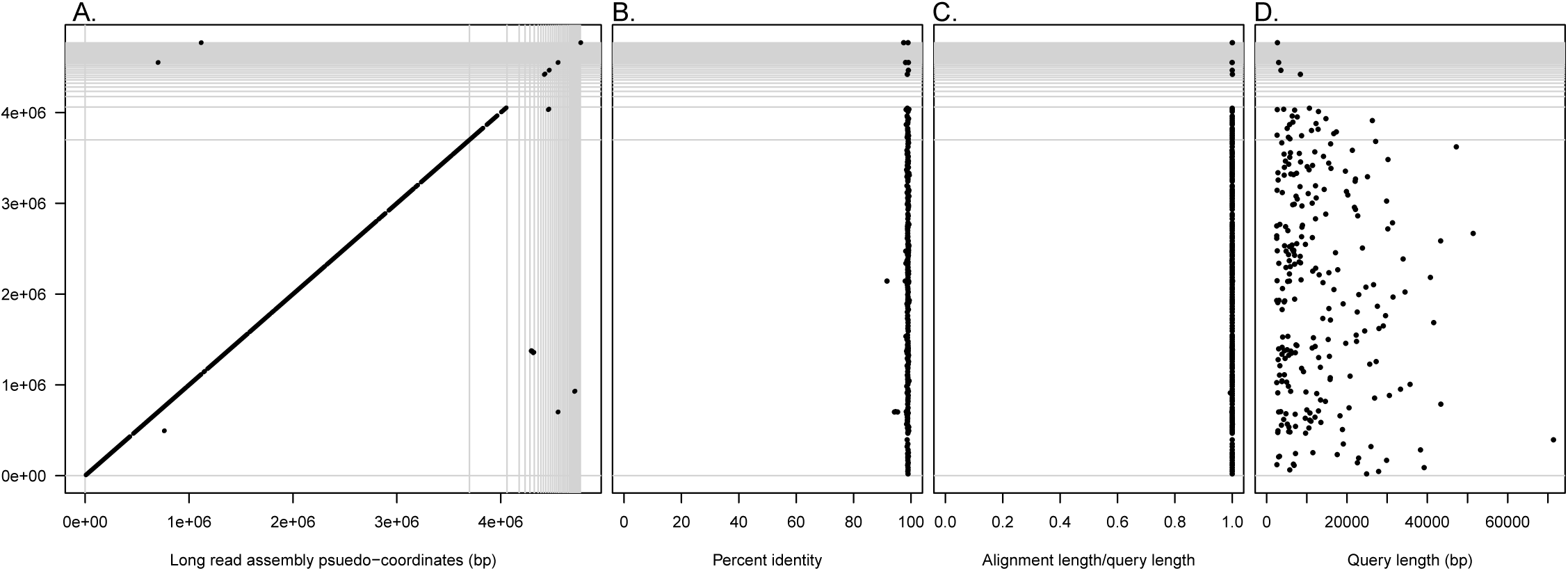
Alignment of SRAC and LRAC sequence for LRAC for SRAC contigs that are **members of AnAOB bin 2**, using **only full length alignments** (*al*2*ql* > 95%). *A).* Dot–plot showing LRAC assembly in pseudo–coordinates, ordered by LRAC contig length in increasing order. Vertical and horizontal lines delineate individual contigs. The first, two LRAC contigs are `tig00000001` and `tig00000002`. Black segments denote individual alignments; *B*) percent identity statistics for aligned sequences. Magnitude of percent identity (*pident*) is plotted on *x*–axis with alignment location on the LRAC assembly plotted on the *y*-axis; *C*) query length statistics for aligned sequences. Magnitude of query length is plotted on *x*–axis with alignment location on the LRAC assembly plotted on the *y*-axis.

We observed that the second longest contig in the assembly, hereafter referred to as `tig00000002` (length: 362k bp; mean base coverage of 42, s.d: 12, see Fig. S2B), showed a similar pattern of alignments from the bin 2 SRAC sequence, suggesting that it is another, shorter fragment of the *Ca.* Brocadia captured in `tig00000001`. The lack of repetition of SRAC alignments relative to those aligned to `tig00000001` suggests this is a distinct genome fragment, not a shorter sub-fragment of the `tig00000001` sequence.

### Refining short read bin composition reduces alignment gaps in long read assembly

Despite the spatially uniform tiling of SRAC sequences from bin 2 sequences across the long read assembly, the dot plots suggest the presence of a number of gaps where no alignments exist. We quantified the extent of this and observed that 21.3% and 44.6% of `tig00000001` and `tig00000002` are not covered by bin 2 SRAC alignments, respectively. Examination of the default bin assignments from MetaBAT (Fig. 1) suggest both bin 2 and bin 6 contain a proportion of outliers, which are be unlikely to derive from their cognate genomes. Because neither the SRAC bin–genomes nor the LRAC sequences can be considered a gold–standard assembly of given member species, we refined the short assembly bin membership and re-examined the extent of tiling across the cognate LRAC sequence, to determine if the occurrence of alignment gaps was determined largely by bin membership, incomplete short read assembly and/or subsequences in the long read assembly associated with sequencing or assembly errors. To expand the total amount of candidate sequence derived from the bin 2 *Ca.* Brocadia genome we included all SRAC contigs at least 1kbp in length and performed the following procedure. We started with the bin 2 contig that contained the full length 16S sequence (contig00593), and extracted contig sets defined by the *k*–nearest neighbours of that contig in the binning plane. We grew these neighbourhoods in increments of 10 contigs, starting with 10 contigs and finishing with 550 contigs (Fig. 10A). For each of these neighbourhoods, we calculated *1)* `CheckM` genome quality statistics (Fig. 10B); *2)* total sequence length (Fig. 10C); *3)* ORF–level annotation statistics summarised at neighbourhood level for family and genus level for *Ca.* Brocadiaceae (Fig. 10D) and *Ca.* Brocadia (Fig. 10E), respectively; *4)* the proportion of LRAC sequence of `tig00000001` and `tig00000002` that contain no aligned SRAC sequence from bin 2 (Fig. 10F); *4)* the number of bin 2 SRAC sequenced that were unaligned to LRAC sequences (Fig. 10G). When calculating gaps in the alignments to LRAC sequence, we only considered full length SRAC aligned sequences (*i.e.* those with *al*2*ql* > 0.95).

**Figure 10:**
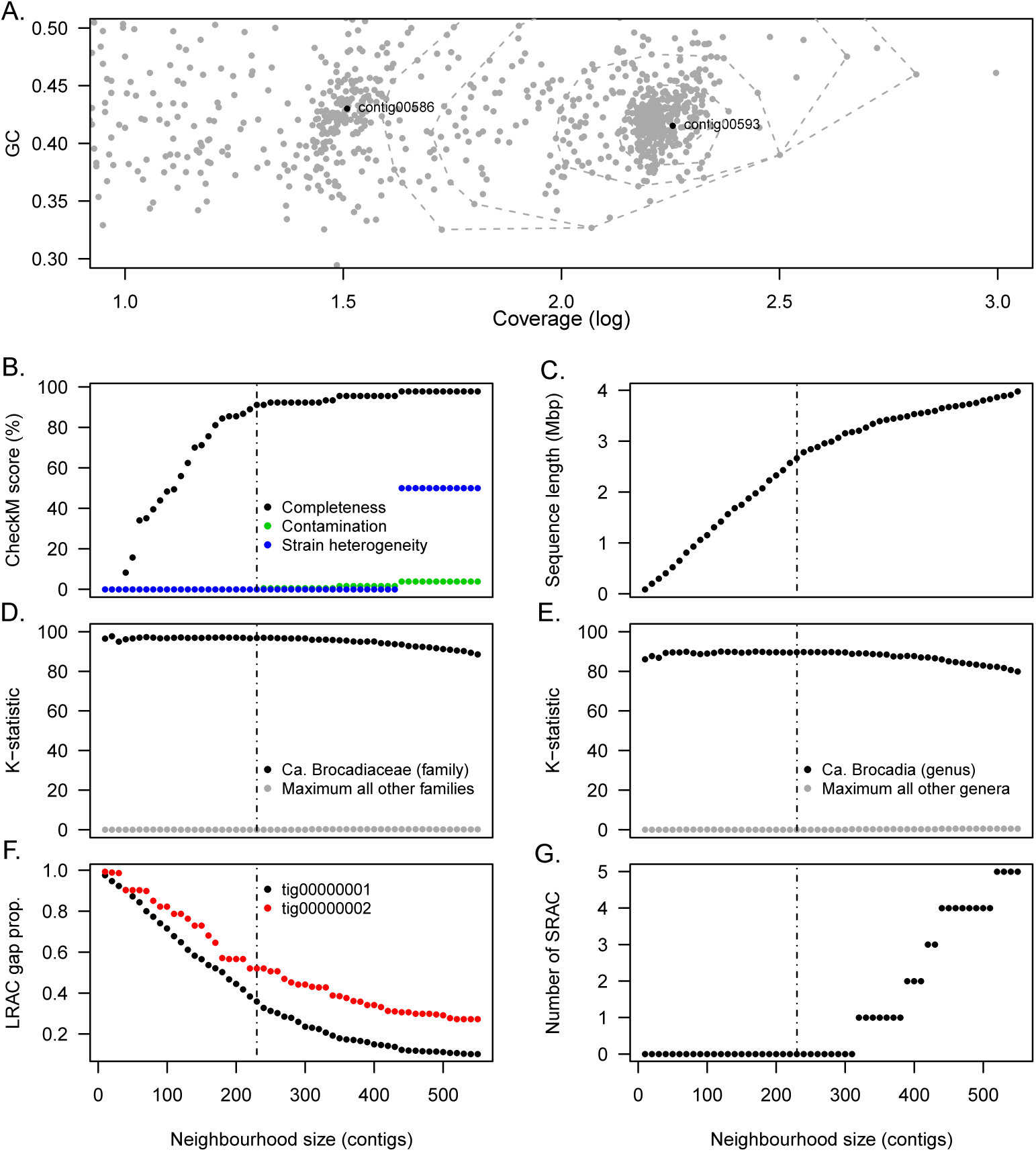
Re–analysis of AnAOB bin 2 using *k*–nearest neighbour contig sets (neighbourhoods) and alignments to LRAC sequence. A). Coverage (*log*_10_)–GC plane showing all contigs wih length ≥ 1*kbp.* Contig neighbourhoods (dashed convex hulls) are defined using the *k*–nearest neighbours of contig00593 which harbours a full length 16S gene annotated to *Ca.* Brocadia. In *A)*–*G*, the value of *k* is plotted on the *x*–axis with the following statistics on the *y*–axis: *B*) `CheckM` genome quality statistics (completeness, contamination and strain heterogeneity); *C)* total sequence length (bp); *D)* ORF–level annotation (*K*) statistics summarised at family level for *Ca.* Brocadiaceae (black) and maximum observed over all other families (grey) *E)* ORF–level annotation (*K*) statistics summarised at genus level for *Ca.* Brocadia (black) and maximum observed over all other genera (grey); *F)* the proportion of LRAC sequence of `tig00000001` and `tig00000002` that contain no aligned SRAC sequence from bin 2; *G)* the number of bin 2 SRAC sequenced that were unaligned to LRAC sequences. When calculating gaps in the alignments to LRAC sequence, we only considered full length SRAC aligned sequences (*i.e.* those with *al*2*ql* > 0.95).

Examining this contig neighbourhood analysis, we observe that 90% completeness as defined by `CheckM` is attained after the *k* = 230 neighbourhood, with non–zero values of contamination and strain heterogeneity appearing at *k* = 240 and *k* = 440, respectively. We note that contamination never exceeds 3.85% which is below the 5% threshold considered to be a high quality draft metagenome assembled genome [17]. Visually, the occurrence of contamination is associated with neighbourhoods starting to impinge on the region of the binning plane associated with contigs in bin 6, as well as outlier contigs from other taxon bins. Total sequence length increased in a near-linear fashion until *k* = 220, attaining a length of 2.67Mb, after which a more gradual increase in length was observed. High values of the *K*-statistic were observed across all neighbourhoods (with minimal signal from other taxa), with a slight decrease towards higher values of *k*. Interestingly, for the *k* = 230 neighbourhood, there remains 36% and 52% of `tig00000001` and `tig00000002` that contain no aligned, full-length SRAC sequence. By *k* = 400 these proportions had dropped to 15% and 33% respectively, with their nadir values of 10% and 27% being observed at *k* = 530 and *k* = 520, respectively. We also note that all SRAC sequences from the neighbourhood are aligned to an LRAC sequence until the *k* = 320 neighbourhood, and across all neighbourhoods a total of 5 SRAC sequences never align to either LRAC sequence. The combination of contamination and strain heterogeneity statistics, along with the appearance of contigs that do not map to LRAC sequences, suggest that the short read derived genome is most complete and uncontaminated between the *k* = 310 and *k* = 430 neighbourhoods, which at the latter contains a total of 3.6Mb of sequence.

Collectively these observations suggest that as defined by LRAC sequences, the AnAOB genome arising from bin 2 of the short read assembly, is just over 4Mb in length and is captured by one long contig (`tig00000001` and one shorter contig (`tig00000002`) of lengths 3,698,683 bp and 362,151 bp, respectively. The gaps in the coverage of SRAC sequences onto these LRAC draft genome is consistent with incomplete assembly of cognate genome using the short read approach. The estimated length of 4Mb is consistent with previous AnAOB genome estimates, including a recent complete, closed assembly of *Kuenenia stuttgartiensis* using PacBio sequencing [5]. Previous short read assemblies, obtained from metagenome survey (MAG), are generally shorter than the estimated length by up to 1Mb, consistent with the above findings.

## Summary and Conclusions

In this paper we examine the capacity of long read data, generated by a Nanopore MinION sequencer, to capture member genomes of a microbial community of low–medium complexity. We show that we can recover a complete, near closed genome of the most abundant community member, specifically an anaerobic ammonium oxidising bacterium (AnAOB) belonging to genus *Ca.* Brocadia. We also show that we can unambiguously resolve the genome of this species without contamination from a lower abundance community member of the same genus. Our results also cast light on the limitations of short read metagenome assembly, and the general challenges associated with accurate recovery of member genomes from complex microbial communities.

Long read metagenomics will likely represent an area of substantive activity over the next several years, particularly in regard to the recovery of complete genomes from member species. Despite great progress in the last decade in developing procedures to obtain short read metagenome–assembled genomes (MAG) [16, 17], substantial limitations of these approaches have become evident, including problems related to the use of multi–sample co-assemblies [38, 39], the challenges of resolving genomes to strain level [40], and the challenges of extracting MAGs from communities of moderate to high ecological complexity [41, 42, 43]. Accordingly, several recent papers have combined short read with emerging complementary techniques such as HiC metagenomics [42, 44], and it is likely to that the combination of short and long read metagenomics will also yield considerable progress in defining higher quality MAGs than is possible using short read metagenome assembly procedures alone. In the present paper, we seek to establish the feasibility of such hybrid long–short metagenomic approaches, starting with communities of low–moderate complexity.

Our analysis is based on the use of a single DNA aliquot co-assayed with both long and short sequencing, thus avoiding the introduction of any difference between the two sets of sequence that could be attributed to ecogenomic differences. Our analysis proceeds on the basis that neither SRAC nor LRAC sequence is known to provide a true reference genome, and so we seek to understand and characterise the degree of agreement between them [45]. Although error prone MinION sequence can be corrected using higher quality short read sequences, in this case, we have deliberately kept the two sources of data separate so as to not introduce any positive bias in our SRAC to LRAC alignment analysis. One consequence of this choice is that direct use of the LRAC assembly is limited in relation to prediction of genes (open reading frames), due to the greatly increased frame–shift rate in coding sequence [13]. Thus in this setting, we would consider our LRAC assembly to be more akin to a genome–wide scaffold for higher quality short read genome fragments, rather than an assembly *per se.* We note that we and others are actively working on correction procedures for frame–shift errors at the time of writing, however, the issues of high error rates in MinION sequence remains a substantial weakness of this technology.

We observe that most of the extant long read assembler or assembly workflows provide broadly comparable results in relation to contig generation (Fig. 2 and Table 2) and recruitment of bin–specific SRAC sequence (Fig. 3), although we note that some behaviours are counter-intuitive. For example, in using `Canu`, increasing the `genomeSize` parameter to ≥ 5Mb results in splitting of the large contig observed in the 3.4Mb version, despite the increased amount of sequence being assembled. We also note that less than 50% of long reads are typically used in the assembly, for reasons that are unclear. Thus, further development of metagenome assemblers and assembly workloads tailored for long read data, or hybrid, short–long read data, is clearly needed. These data were obtained during a phase of rapid development of MinION kits and protocols, only generating under 1Gb of sequence from these complex environmental samples, and we anticipate more comprehensive results using more recent datasets, given the improvement in yield, read length and quality now available from the same technology.

The concept of genome completeness remains problematic in MAG–based analyses of shotgun metagenome data. Current methods for estimating genome completeness from short read data are based on marker gene sets whose membership is defined by the fact of having a single genomic copy. Typically, these are of order of around 100 genes, and by definition such sets cannot accurately estimate the completeness of a genome. In the present case, we see that high level completeness in the SRAC sequence are occurring at total sequence length that is around 1.4Mb less than the genome length inferred from LRAC based analyses. In fact we would argue that given the complexity of how short read contigs from many member species will be distributed in a given binning space, there must remain a fundamental ambiguity in resolving genome content using these approaches, as highlighted by our neighbourhood based reanalysis of bin 2. While the backbone of the genome is certainly captured under these criteria, knowing when one has accurately captured a whole genome remains challenging and reinforces the view that MAG–based analysis can likely only resolve a working model of the genome (or pangenome) of a given community member in most circumstances. In the present study we have also used completely automated methods, with no recourse to manual or subjective decision making that appear to be commonly deployed in the refinement of MAG draft genomes and represent a challenge to the conduct of reproducible research in this area.

Further development of long read metagenomics may in fact obviate the need for complex binning procedures, with methods refocusing on the evaluation of LRAC sequences as putative draft genomes. Either way, it would appear as if conduct of hybrid short–long read metagenome surveys will be essential for resolving these challenges for the foreseeable future. The engagement of other complementary approaches, namely single cell metagenomics [46], HiC–metagenomics [44] or FACS–generated mini–metagenomes [43] in combination with MAG approaches, will likely permit substantial progress.

## List of Supplementary Materials

Supplementary items appear with this document unless otherwise stated.

- Table S1: Summary statistics for the short read assembly
- Table S2: CheckM statistics for short read assembly bins
- Table S3: *K*–statistics for short read assembly bins at phylum, family and genera levels
- Table S4: Summary of phyla–level analysis of long read data using MEGAN–LR
- Table S5: Summary of genera–level analysis of long read data using MEGAN–LR
- Supplementary Datafile 1: Alignment statistics for SRAC derived 16S (xlsx file)
- Figure S1: Per–base long read coverage for AnAOB–associated LRAC sequences
- Figure S2: Relationship between length and coverage in the short–read metagenome assembly

## Author contributions

The study was conceptualised by RBHW and designed by RBHW and IB. GN, YYL and SW setup and operated enrichment reactors, and obtained samples with IB. IB performed DNA extractions and performed long read sequencing, with IB and FL designed long read sequencing experiments. DID performed short read sequencing. KA, RBHW, XHL and DHH designed and/or performed data analysis. All authors contributed to data interpretation. RBHW wrote the manuscript with specific contributions from all other authors.

## Acknowledgements

This research was supported by the Singapore National Research Foundation and Ministry of Education under the Research Centre of Excellence Programme, and by program grant 1301–IRIS–59 from the National Research Foundation (NRF). We thank Gavin Huttley (Australian National University) for critical feedback on sequence analysis. Early version of this work were presented at APBC2018 (Yokohama, Japan, January 15–17 2018) and ISME17 (Leipzig, Germany, August 12–17 2018)

---

**Table S1:** Summary statistics for the short read assembly

| Measure |  |
| --- | --- |
| Contigs ($N$) | 38,988 |
| Total length (bp) | 75,821,912 |
| Longest contig (bp) | 838,453 |
| Shortest contig (bp) | 500 |
| Contigs>500 bp ($N$) | 38,910 |
| Contigs>1000 bp ($N$) | 14,450 |
| Contigs>10k bp ($N$) | 813 |
| Contigs>100k bp ($N$) | 37 |
| Contigs>1M bp ($N$) | 0 |
| Mean length (bp) | 1,945 |
| N50 | 4,145 |

**Table S2:** Summary of phyla–level analysis of long read data using MEGAN–LR

| Taxonomic path (domain, phylum) | Number of alignments | Proportion of total (%) |
| --- | --- | --- |
| Bacteria;PVC group;Planctomycetes | 1.57E+08 | 53.94 |
| Bacteria;Proteobacteria | 6.97E+07 | 23.97 |
| Bacteria;Terrabacteria group;Chloroflexi; | 2.10E+07 | 7.21 |
| Bacteria;Terrabacteria group;Armatimonadetes; | 1.79E+07 | 6.14 |
| Bacteria;FCB group;Bacteroidetes/Chlorobi group;Bacteroidetes; | 1.24E+07 | 4.26 |
| Bacteria;Nitrospirae; | 3.27E+06 | 1.12 |
| Bacteria;Terrabacteria group;Actinobacteria <phylum>; | 2.72E+06 | 0.93 |
| Bacteria;Acidobacteria; | 1.96E+06 | 0.67 |
| Bacteria;Terrabacteria group;Cyanobacteria/Melainabacteria group;Cyanobacteria; | 1.29E+06 | 0.44 |
| Bacteria;FCB group;Bacteroidetes/Chlorobi group;Ignavibacteriae; | 1.25E+06 | 0.43 |
| Bacteria;PVC group;Candidatus Omnitrophica; | 7.34E+05 | 0.25 |
| Bacteria;unclassified Bacteria;unclassified Bacteria (miscellaneous); | 5.99E+05 | 0.21 |
| Bacteria;Terrabacteria group;Firmicutes; | 4.77E+05 | 0.16 |
| Bacteria;FCB group;Bacteroidetes/Chlorobi group;Chlorobi; | 3.99E+05 | 0.14 |
| Eukaryota;Opisthokonta;Metazoa;Eumetazoa; | 1.75E+05 | 0.06 |
| Bacteria;unclassified Bacteria;Bacteria candidate phyla;Patescibacteria group; | 1.75E+05 | 0.06 |

**Table S3:**
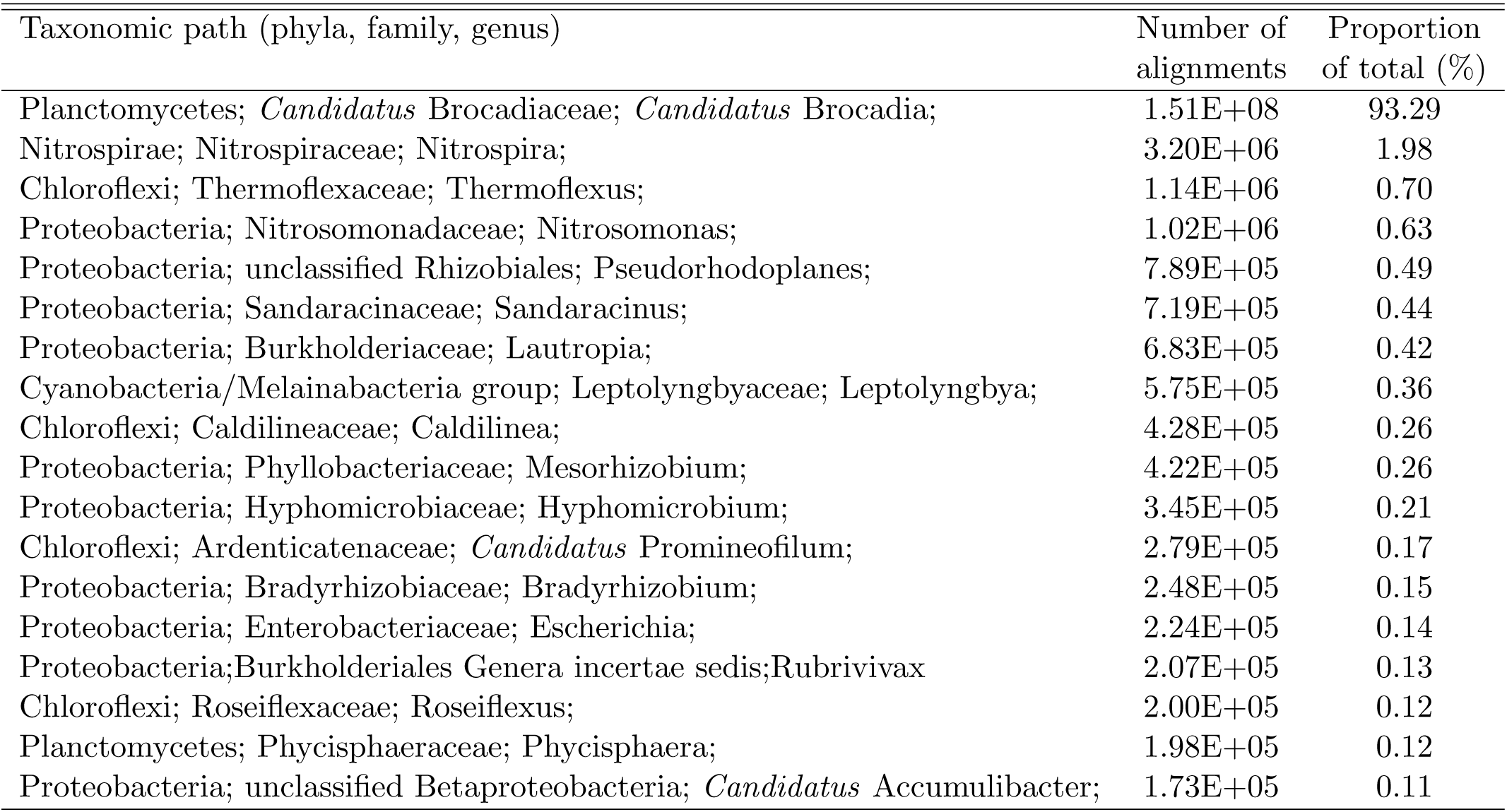
Summary of genera–level analysis of long read data using MEGAN–LR

**Table S4:** CheckM genome quality statistics for short read assembly bins

| Bin | Lineage | Completeness | Contamination | Strain heterogeneity | $\widehat{Cov}$ | $\widehat{GC}$ |
| --- | --- | --- | --- | --- | --- | --- |
| 8 | k__Bacteria (UID203) | 99.69 | 149.37 | 0.93 | 9 | 0.70 |
| 6 | k__Bacteria (UID2565) | 98.84 | 1.65 | 0.00 | 31 | 0.41 |
| 7 | k__Bacteria (UID1452) | 98.18 | 0.00 | 0.00 | 22 | 0.63 |
| 1 | k__Bacteria (UID1452) | 95.83 | 0.93 | 0.00 | 180 | 0.61 |
| 2 | k__Bacteria (UID2565) | 95.40 | 3.85 | 50.00 | 150 | 0.42 |
| 5 | k__Bacteria (UID1452) | 94.55 | 2.27 | 0.00 | 31 | 0.53 |
| 11 | k__Bacteria (UID203) | 85.34 | 26.72 | 28.30 | 9 | 0.68 |
| 4 | k__Bacteria (UID203) | 75.86 | 0.00 | 0.00 | 44 | 0.67 |
| 9 | k__Bacteria (UID2565) | 62.52 | 10.23 | 0.00 | 8 | 0.64 |
| 13 | k__Bacteria (UID1452) | 62.01 | 16.86 | 3.85 | 6 | 0.66 |
| 10 | k__Bacteria (UID1452) | 58.60 | 0.99 | 0.00 | 8 | 0.54 |
| 16 | k__Bacteria (UID203) | 3.26 | 0.00 | 0.00 | 7 | 0.65 |
| 3 | root (UID1) | 0.00 | 0.00 | 0.00 | 142 | 0.39 |
| 15 | root (UID1) | 0.00 | 0.00 | 0.00 | 8 | 0.59 |
| 14 | root (UID1) | 0.00 | 0.00 | 0.00 | 7 | 0.70 |
| 12 | root (UID1) | 0.00 | 0.00 | 0.00 | 7 | 0.65 |

$\widehat{Cov}$ : mean coverage of member contigs
$\widehat{GC}$ : mean GC content of member contigs

**Table S5:**
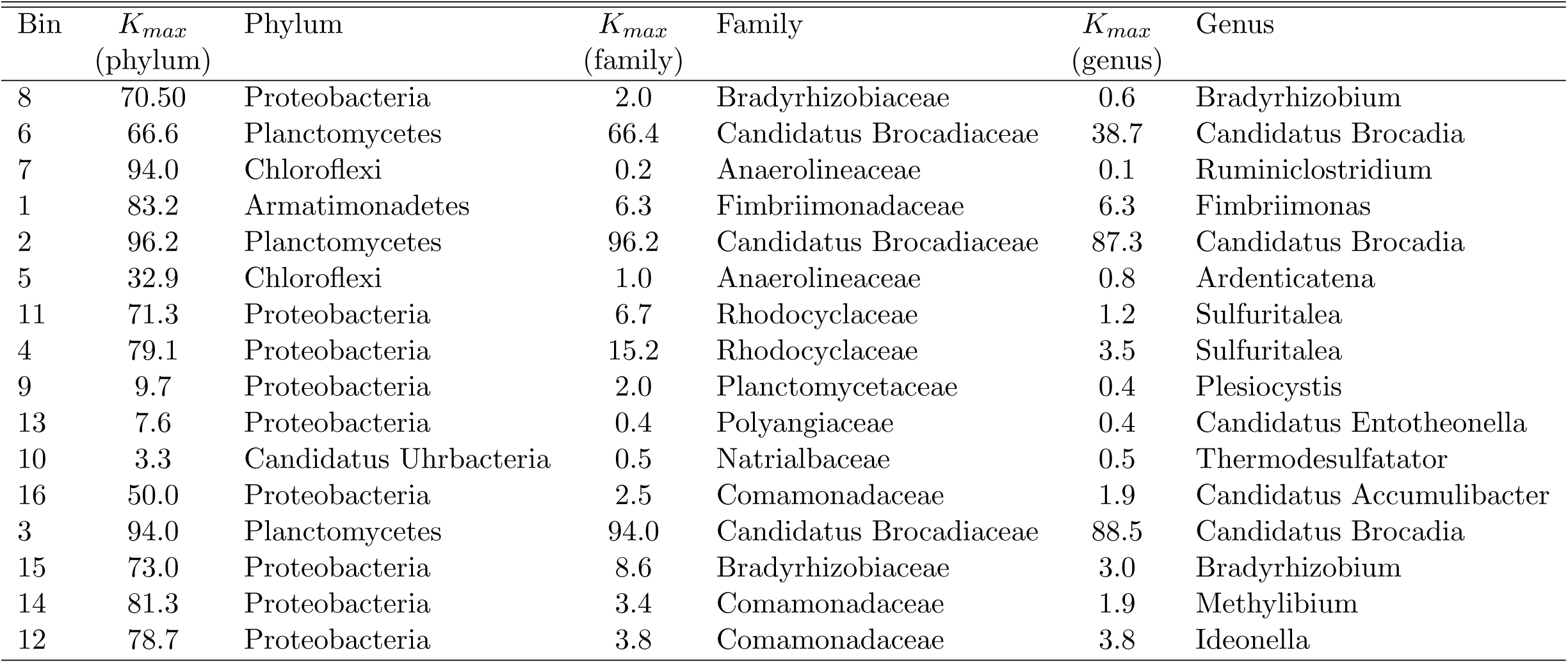
*K*–statistics for short read assembly bins at phylum, family and genera levels

**Figure S1:**
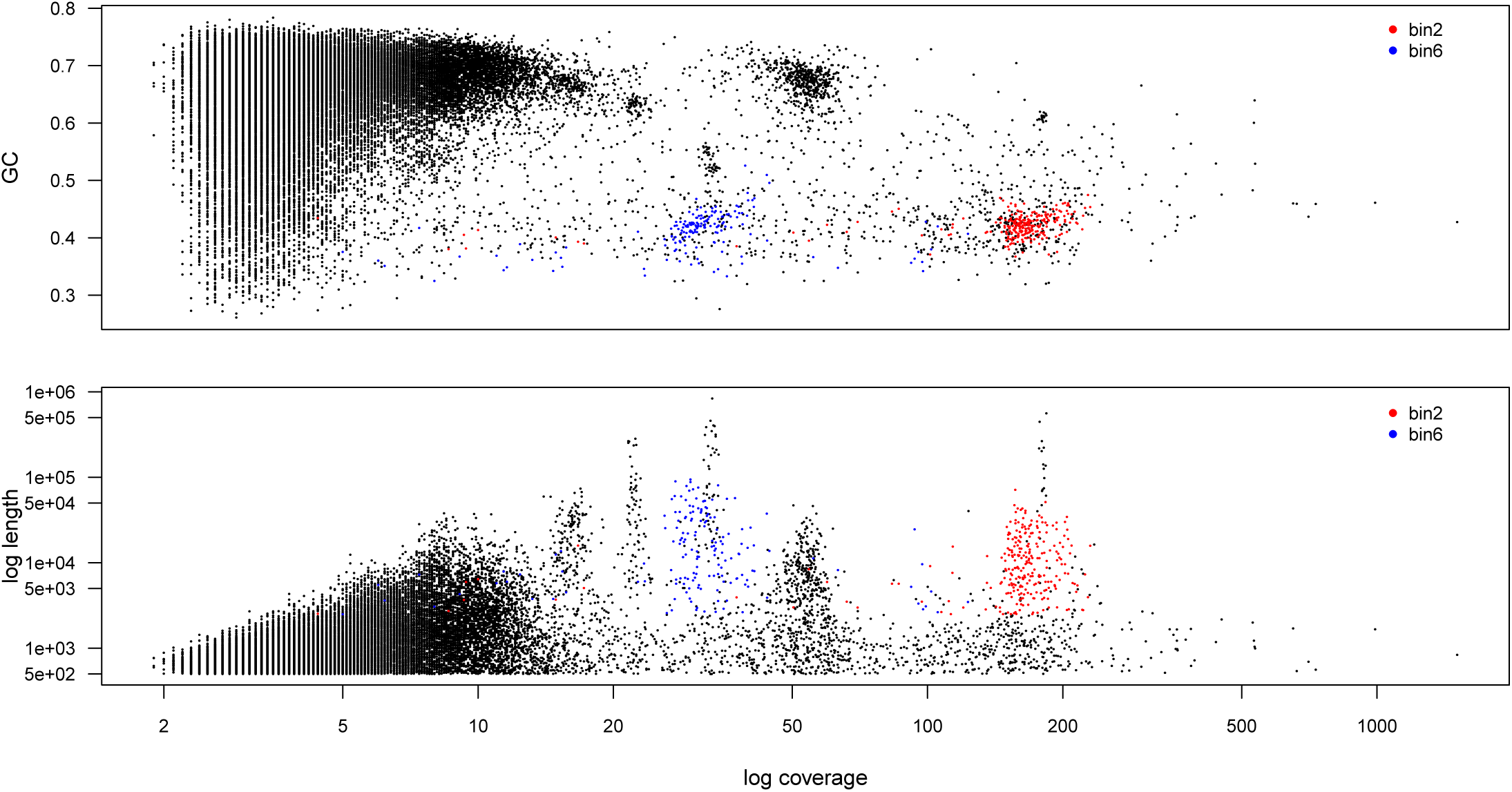
Inter-relationship between contig GC content, coverage and length in the short read assembly. Each datapoint is a contig. *Upper panel*: GC–proportion plotted against log coverage; *Lower panel*: Log length plotted against log coverage. In both plots, contigs that are members of bin 2 are highlighted in red and contigs that are members of bin 6 are highlighted in blue.

**Figure S2:**
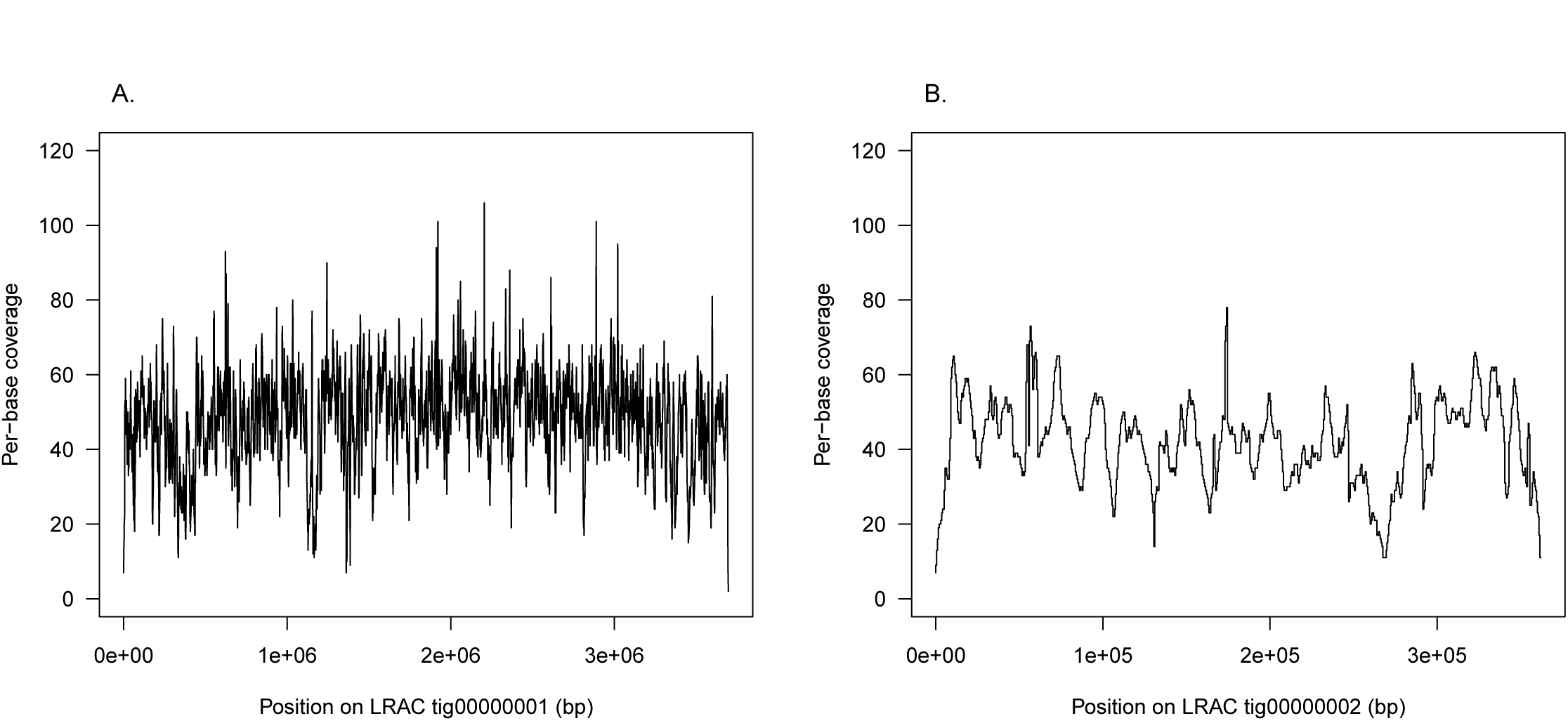
Smoothed per-base long read coverage for LRAC sequences *A* `tig00000001`; and *B*: `tig00000002`. A running median has been applied to the raw per-base coverage data using a 1001 bp length window, and then the smoothed data has been subsampled at an interval of 500 basepairs

## References

[1] Loman, N.J., Quick, J., Simpson, J.T. (2015) A complete bacterial genome assembled *de novo* using only Nanopore sequencing data. Nat. Methods 12 (8): 733–5

[2] Wick, R.R., Judd, L.M., Gorrie, C.L., Holt, K.E. (2017). Completing bacterial genome assemblies with multiplex MinION sequencing, Microb. Genom. 3(10): e000132

[3] Doyle, L.E., Williams, R.B.H., Rice, S.A., Marsili, E., Lauro, F.M. (2018). Draft genome sequence of *Enterobacter* sp. Strain EA–1, an electrochemically active microorganism isolated from tropical sediment, Genome Announcements 6(9): e00111–18.

[4] Daebeler, A., Herbold C.W., Vierheilig, J., Sedlacek C.J., Pjevac, P., Albertsen, M., Kirkegaard, R.H., de la Torre, J.R., Daims, H., Wagner, M. (2018). Cultivation and genomic analysis of “*Candidatus* Nitrosocaldus islandicus”, an obligately thermophilic, ammonia–oxidizing Thaumarchaeon from a hot spring biofilm in Graendalur Valley, Iceland. Frontiers in Microbiology 9: 193.

[5] Frank, J., Lücker, S., Vossen, R.H.A.M., Jetten, M.S.M., Hall, R.J., Op den Camp, H.J.M., Anvar, S.Y. (2018). Resolving the complete genome of *Kuenenia stuttgartiensis* from a membrane bioreactor enrichment using Single–Molecule Real–Time sequencing. Scientific Reports. 8(1): 4580.

[6] Andersen, M.H., McIlroy, S.J., Nierychlo, M., Nielsen, P.H., Albertsen, M. (2018). Genomic insights into *Candidatus* Amarolinea aalborgensis gen. nov., sp. nov., associated with settleability problems in wastewater treatment plants, Systematic and Applied Microbiology, available online 16 August 2018 https://doi.org/10.1016/j.syapm.2018.08.001

[7] Driscoll, C.B., Otten T.G., Brown, N.B., Dreher, T.W. (2017). Towards long-read metagenomics: complete assembly of three novel genomes from bacteria dependent on a diazotrophic cyanobacterium in a freshwater lake co–culture, Standards in Genomic Sciences. 12: 9

[8] Slaby, B.M., Hackl, T., Horn, H., Bayer, K., Hentschel, U. (2017). Metagenomic binning of a marine sponge microbiome reveals unity in defense but metabolic specialization, ISME Journal, 11: 2465–2478.

[9] Frank, J.A., Pan, Y., Tooming–Klunderud, A., Eijsink, V.G.H., McHardy, A.C., Nederbragt, A.J. (2016). Improved metagenome assemblies and taxonomic binning using long–read circular consensus sequence data, Scientific Reports, 6: 25373.

[10] Singer, E., Andreopoulos, B., Bowers, R.M., Lee, J., Deshpande, S., Chiniquy, J., Ciobanu, D., Klenk, H-P., Zane, M., Daum, C., Clum, A., Cheng, J-F., Copeland, A., Woyke, T. (2016) Next generation sequencing data of a defined microbial mock community. Scientific Data 3:160081.

[11] Brown, B.L., Watson, M., Minot, S.S., Rivera, M.C. Franklin, R.B. (2017). MinION nanopore sequencing of environmental metagenomes: a synthetic approach. GigaScience 6: 1–10.

[12] Nanopore GridION and PromethION Mock Microbial Community Data Community Release, Release 2 (2018-10-17). https://github.com/LomanLab/mockcommunity

[13] Huson, D.H., Albrecht, B., Bagci, C., Bessarab, I., Gorska, A., Jolic, D., Williams, R.B.H. (2018). MEGAN–LR: New algorithms allow accurate binning and easy interactive exploration of metagenomic long reads and contigs, Biology Direct 13: 6

[14] Laczny, C.C., Kiefer, C., Galata, V., Fehlmann, T., Backes, C., Keller, A. (2017). BusyBee Web: metagenomic data analysis by bootstrapped supervised binning and annotation. Nucleic Acids Res. 45 (W1): W171–W179

[15] Liu, X.H., Arumugam, K., Natagaran, G., Seviour, T.W., Drautz-Moses, D.I., Wuertz, S., Law, Y.Y., Williams, R.B.H. (2018). Draft genome sequence of an “*Candidatus* Brocadia” bacterium enriched from tropical-climate activated sludge, Genome Announcements, in press. Preprint biorxiv.org/content/early/2017/04/24/123943

[16] Quince, C., Walker, A.W., Simpson, J.T., Loman, N.J., Segata, N. (2017), Shotgun metagenomics, from sampling to analysis, Nature Biotechnology 35: 833–84.

[17] Bowers, R.M., Kyrpides, N.C. Stepanauskas, R., Harmon–Smith, M., Doud, D. et al. (2017). Minimum information about a single amplified genome (MISAG) and a metagenome–assembled genome (MIMAG) of bacteria and archaea, Nature Biotechnology 35: 725–731.

[18] Koren, S., Walenz, B.P., Berlin, K., Miller, J.R., Bergman, N.H., Phillippy, A.M. (2017). Canu: scalable and accurate long–read assembly via adaptive *k*–mer weighting and repeat separation, Genome Research, 27(5): 722–736.

[19] Li, H. (2016). Minimap and miniasm: fast mapping and de novo assembly for noisy long sequences, Bioinformatics 32(14): 2103–2110.

[20] SMARTdenovo https://github.com/ruanjue/smartdenovo

[21] Recanati, A., Brüls, T., d’Aspremont, A. (2017). A spectral algorithm for fast *de novo* layout of uncorrected long nanopore reads, Bioinformatics 33(20): 3188–3194.

[22] Wick, R.R., Judd, L.M., Gorrie, C.L. Holt, K.E. (2017). Unicycler: Resolving bacterial genome assemblies from short and long sequencing reads, PLoS Computational Biology 13(6): e1005595.

[23] Altschul, S.F., Gish, W., Miller, W., Myers, E.W., Lipman, D.J. (1990). Basic local alignment search tool. J. Mol. Biol. 215: 403–410.

[24] O’Leary NA, Wright MW, Brister JR, Ciufo S, Haddad D et al. (2016). Reference sequence (RefSeq) database at NCBI: current status, taxonomic expansion, and functional annotation. Nucleic Acids Res. 44(D1): D733–45

[25] Martin, M. (2011). Cutadapt removes adapter sequences from high–throughput sequencing reads. EMBnet.journal, 17(1): 10–12.

[26] Xie, C., Goi, W.C., Huson, D.H., Little, P.F.R., Williams, R.B.H. (2016). RiboTagger: fast and unbiased 16S/18S profiling using whole community shotgun metagenomic or metatranscriptome surveys, BMC Bioinformatics 17(Suppl 19): 1378

[27] Pruesse, E., Quast, C., Knittel, K., Fuchs, B.M., Ludwig, W., Peplies, J., Glckner, F.O. (2007). SILVA: a comprehensive online resource for quality checked and aligned ribosomal RNA sequence data compatible with ARB. Nucleic Acids Research 35: 7188–7196.

[28] Edgar, R.C. (2017). SEARCH_16S: A new algorithm for identifying 16S ribosomal RNA genes in contigs and chromosomes. http://biorxiv.org/content/early/2017/04/04/124131

[29] Pruesse, E., Peplies, J., Glöckner, F.O. (2012) SINA: accurate high–throughput multiple sequence alignment of ribosomal RNA genes. Bioinformatics 28: 1823–1829

[30] Zhu, W., Lomsadze, A., Borodovsky, M. (2010). *Ab initio* gene identification in metagenomic sequences. Nucleic Acids Research 38: e132.

[31] Buchfink, B., Xie, C., Huson, D.H. (2015). Fast and sensitive protein alignment using DIAMOND, Nature Methods 12(1): 59–60.

[32] Kang, D.D., Froula, J., Egan, R., Wang, Z. (2015). MetaBAT, an efficient tool for accurately reconstructing single genomes from complex microbial communities, PeerJ, 3, e1165.

[33] Parks, D.H., Imelfort, M., Skennerton, C.T., Hugenholtz, P., Tyson, G.W. (2015). CheckM: assessing the quality of microbial genomes recovered from isolates, single cells, and metagenomes” Genome Research, 25, 1043–1055.

[34] Williams, R.B.H., Liu, X.H., Arumugam, K. (2018). Annotation statistics for exploratory analysis of metagenome assembly binning data, unpublished manuscript.

[35] Porechop. https://github.com/rrwick/Porechop

[36] Li, H. (2018). Minimap2: pairwise alignment for nucleotide sequences. Bioinformatics 34 (18): 3094–3100.

[37] Li, H., Handsaker, B., Wysoker, A., Fennell, T., Ruan, J., Homer, N., Marth, G., Abecasis, G., Durbin, R.; 1000 Genome Project Data Processing Subgroup. The Sequence Alignment/Map format and SAMtools, Bioinformatics. 25(16): 2078–2079.

[38] Olm, M.R., Brown, C.T., Brooks, B., Banfield, J.F. (2018). dRep: a tool for fast and accurate genomic comparisons that enables improved genome recovery from metagenomes through de–replication, ISME J. 11(12): 2864–2868

[39] Miller, I.J., Rees, E.R., Ross, J., Miller, I., Baxa, J., Lopera, J., Kerby, R.L., Rey, F.E., Kwan, J.C. (2018). Autometa: Automated extraction of microbial genomes from individual shotgun metagenomes, bioRxiv 251462; doi: https://doi.org/10.1101/251462.

[40] Quince, C., Delmont, T.O., Raguideau, S., Alneberg, J., Darling, A.E., Collins, G., Eren, A.M. (2017). DESMAN: a new tool for de novo extraction of strains from metagenomes, Genome Biol. 18(1): 181

[41] Delmont, D.O., Quince, C., Shaiber, A., Esen, O.E., Lee, S.T.M., Rappé, M.S., McLellan, S.L., Liicker, S., Eren, A.M. Nitrogen–fixing populations of Planctomycetes and Proteobacteria are abundant in surface ocean metagenomes, Nature Microbiology 3: 804–813.

[42] Stewart, R.D., Auffret, M.D., Warr, A., Wiser, A.H., Press, M.O., Langford, K.W., Liachko, I., Snelling, T.J., Dewhurst, R.J., Walker, A.W., Roehe, R., Watson, M. (2018). Assembly of 913 microbial genomes from metagenomic sequencing of the cow rumen, Nature Communications, 9: 870

[43] Ji, P., Zhang, Y.M., Wang, J.F., Zhao, F.Q. (2017). MetaSort untangles metagenome assembly by reducing microbial community complexity. Nature Communications, 8, 14306.

[44] Burton, J.N., Liachko, I., Dunham, M.J., Shendure, J. (2014). Species–level deconvolution of metagenome assemblies with Hi–C–based contact probability maps. G3 (Bethesda), 4(7): 1339–46

[45] Bland, J.M., Altman, D.G. (1986). Statistical methods for assessing agreement between two methods of clinical measurement. Lancet 327 (8476): 307–10.

[46] Lan, F., Demaree, B., Ahmed, N., Abate, A.R. (2017). Single-cell genome sequencing at ultra-high-throughput with microfluidic droplet barcoding, Nat. Biotechnol. 35(7): 640–646.

